# Reg4 defenses the intestine against *Salmonella* infection via binding the flagellin

**DOI:** 10.1101/2021.06.03.447020

**Authors:** Weipeng Wang, Ying Lu, Xinbei Tian, Shanshan Chen, Jun Du, Wei Cai, Yongtao Xiao

## Abstract

*Salmonella* Typhimurium is gram-negative flagellated bacteria that can cause food-borne gastroenteritis and diarrhea in humans and animals. The regenerating islet-derived family member 4 (Reg4) is overexpressed in the gastrointestinal tract during intestinal inflammation. However, the role of Reg4 in the intestinal inflammation induced by *Salmonella* Typhimurium is largely unknown. In this study, we reported for the first time that Reg4 has bactericidal activity against intestinal infection caused by *Salmonella* Typhimurium. In vivo, Reg4 could reduce the colonization of *Salmonella* Typhimurium and attenuate intestinal inflammation in the *Salmonella* Typhimurium-infected model. Additionally, the mice with the epithelial cell specific deletion of *Reg4* (*Reg4^ΔIEC^*) exhibited more severe intestinal inflammation and more colonization of *Salmonella* Typhimurium. However, the administration of Reg4 could reverse these negative impacts. In vitro, Reg4 protein was showed to inhibit the growth of *Salmonella* Typhimurium. We further investigate the function motif of Reg4 and find that the “HDPQK” motif in Reg4 is essential to its bactericidal activity. Reg4 exerted the bactericidal effect by binding to the flagellin of *Salmonella* Typhimurium and suppressing its motility, adhesion, and invasion to the intestinal epithelia. In conclusion, our findings identify Reg4 as a novel antimicrobial peptide against infection by *Salmonella* Typhimurium and explore its possible mechanism, which may be of great significance for developing novel agents against flagellated micro pathogens.

## Introduction

*Salmonella enterica serovar* Typhimurium (*Salmonella* Typhimurium) is a common zoonotic enteropathogenic bacteria that can cause enterocolitis and diarrhea, which is the leading cause of morbidity and mortality in people worldwide. Young children and immunosuppressed patients are at risk of systemic dissemination of this pathogen and require antimicrobial treatment(1). Unfortunately, over the years, the use of these antimicrobial agents has promoted the emergence of resistant strains(2, 3). The resistance to multiple antimicrobial agents resulted in increased treatment failure rates of *Salmonella* Typhimurium infection (>55%), leading to a global public health crisis(1). Therefore, the need to develop efficient strategies to combat antibiotic resistance is urgent.

Novel antimicrobial peptides (AMPs) have already evolved into formidable candidates for antibiotic substitute materials against pathogenic infections. In the intestine, AMPs derived from epithelial cells are essential to avoid colonization and invasion of opportunistic pathogens and maintain immune homeostasis (4). Regenerating islet-derived (Reg) proteins have emerged as multifunctional agents with bactericidal properties, pro-proliferative, differentiation-inducing, and anti-apoptotic(5). The Reg family comprises four groups (Reg1, Reg2, Reg3, and Reg4) of proteins based on their primary structure(5). REG4 was recently discovered and characterized from the cDNA library of ulcerative colitis(6). Reg4 is physiologically expressed in the enteroendocrine cells of rodents and humans and expands into epithelial cells during intestinal inflammation(7–9). The Reg4 mRNA was highly expressed in the inflamed epithelium of the gastrointestinal tract in patients with ulcerative colitis and Crohn’s disease(10, 11). Although Reg4 exhibits the lowest similarity with any of the other Reg proteins, it is also involved in the lectin-related biological processes by containing a sequence motif homologous to calcium-dependent (C-type) lectin-like domain(5). These suggests that Reg4 may play an important role in the progression of Intestinal inflammation. Surprisingly though, to our knowledge, very little is known about the role of Reg4 during *Salmonella* Typhimurium infection.

In this study, we purified the recombinant human REG4 and mouse Reg4 proteins and investigate the exact role of Reg4 in intestinal inflammation using the *Salmonella* Typhimurium-infected mice and *ex vivo* experiments. In addition, we further determined the functional motif of Reg4 and clarified how Reg4 protein exerts its bactericidal activity against Salmonella.

## Results

### Reg4 is highly expressed in intestinal mucosa and increasingly expressed during inflammation

As shown in Fig. 1A, *Reg4* was mainly expressed in the mouse intestinal mucosa. The expression of *Reg4* mRNA was highest in the cecum and then in the colon and small intestine (Fig. 1A). The gene expression dataset GSE23063 and GSE27000 was downloaded from the GEO database and analyzed to determine the expression of Reg4 during *Salmonella* infection. As presented in Fig.1B, *Salmonella* infection induced the higher expression of Reg4 in the porcine intestinal epithelial cell lines. When infected with *Salmonella*, the pigs with higher expression of *Reg4* had lower shedding of *Salmonella* compared to the pigs with low expression of *Reg4* (Fig. 1C). Consistently, in this study, the real-time PCR (RT-PCR) showed the higher expression of *Reg4* mRNA in the colonic mucosa of mice infected with *Salmonella* (Fig. 1D). Consistent with the findings in *Reg4* mRNA, Immunofluorescence (IF) showed that that Reg4 protein exclusively expressed in intestinal mucosa (Fig. 1E).

**Figure 1.**
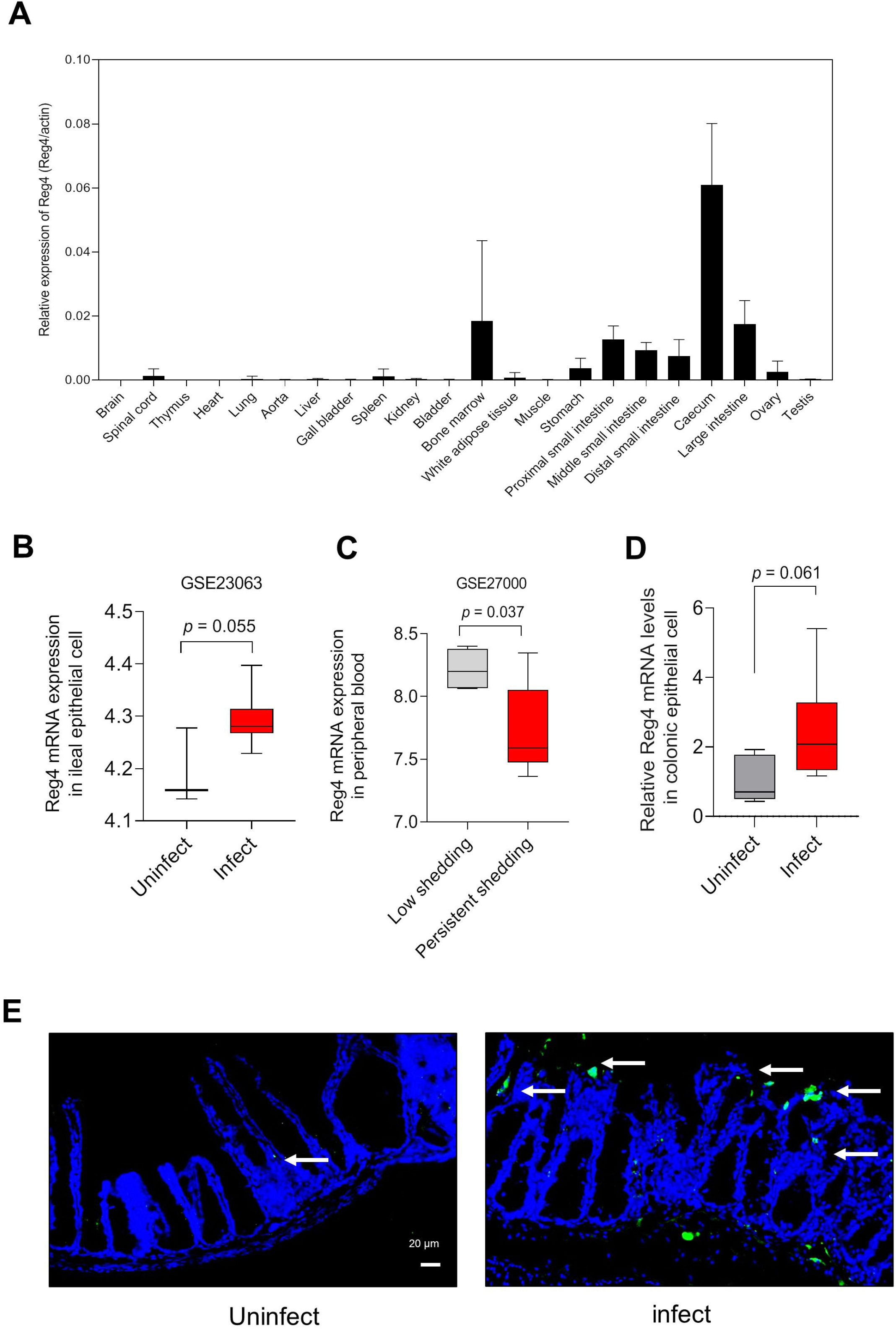
*Reg4* is expressed in intestine and aberrant expressed during *Salmonella*-induced inflammation. **(A)** Expression of *Reg4* mRNA in normal tissues of mice. **(B)** Expression of *Reg4* mRNA in ileum epithelial cells of *Salmonella*-infected pig was analyzed using the GEO database (GSE23063). Unpaired t test was used for statistical analysis. **(C)** Expression of *Reg4* mRNA in peripheral blood of *Salmonella*-infected pig was analyzed using the GEO database (GSE27000). Unpaired t test was used for statistical analysis. **(D)** Expression of *Reg4* mRNA in the mucosa of mice colon (n = 6 per group). Unpaired t test was used for statistical analysis. Unpaired t test was used for statistical analysis; ns, not significant (*p* ≥ 0.05); * *p* < 0.05; ** *p* < 0.01, *** *p* < 0.005; **** *p* < 0.001. **(E)** Immunofluorescence images of mouse cecum stained for Reg4 (green) and DAPI (blue). White arrows highlight Reg4.

### Reg4 is protective against *Salmonella* Typhimurium infection

To test whether recombinant Reg4 has bactericidal activity against with *Salmonella* Typhimurium infection, we firstly purified the recombinant mouse Reg4 and human REG4 proteins (Fig. 2A-B) and incubated them with *Salmonella* Typhimurium. We found that the growth of *S. Enteritidis* was suppressed by the human Reg4 protein in a dose-dependent manner (Fig. 3A-B). Similarly, mouse Reg4 protein significantly inhibited the growth of *S. Tm* (Fig. 3A-B). *In vivo*, Salmonella infection induce the decease of body weight and intestinal lesion, and this damage could be significantly reduced by administration of the recombinant mouse Reg4 (Fig. 3C-E). The histological score of cecum was evidently improved when the infected mice was administrated with Reg4 protein (Fig. 3D-E). For chronic infection, the Reg4 improved the survival rate of the infected mice (Fig. 3F). Additionally, compared with these untreated infected mice, the inflammatory cytokine levels in the serum and colonic mucosa of the treated infected mice were decreased (Fig. 4A-C). To further investigate the possible protective role of Reg4 in the intestinal infection induced by *Salmonella* Typhimurium, we generated mice with an epithelial cell specific deletion of Reg4 (*Reg4^ΔIEC^*) as reported previously(12). It was found that *Reg4^ΔIEC^* mice were more sensitive to *S. Tm* infection than wild-type mice and exhibited more weight loss (Fig.3C), higher histological score (Fig.3D-E), and higher levels of the inflammatory cytokines IFN-γ, and IL-22 in serum (Fig. 4A-B) and expression of inflammatory genes in colonic mucosa (Fig. 4C).

**Figure 2.**
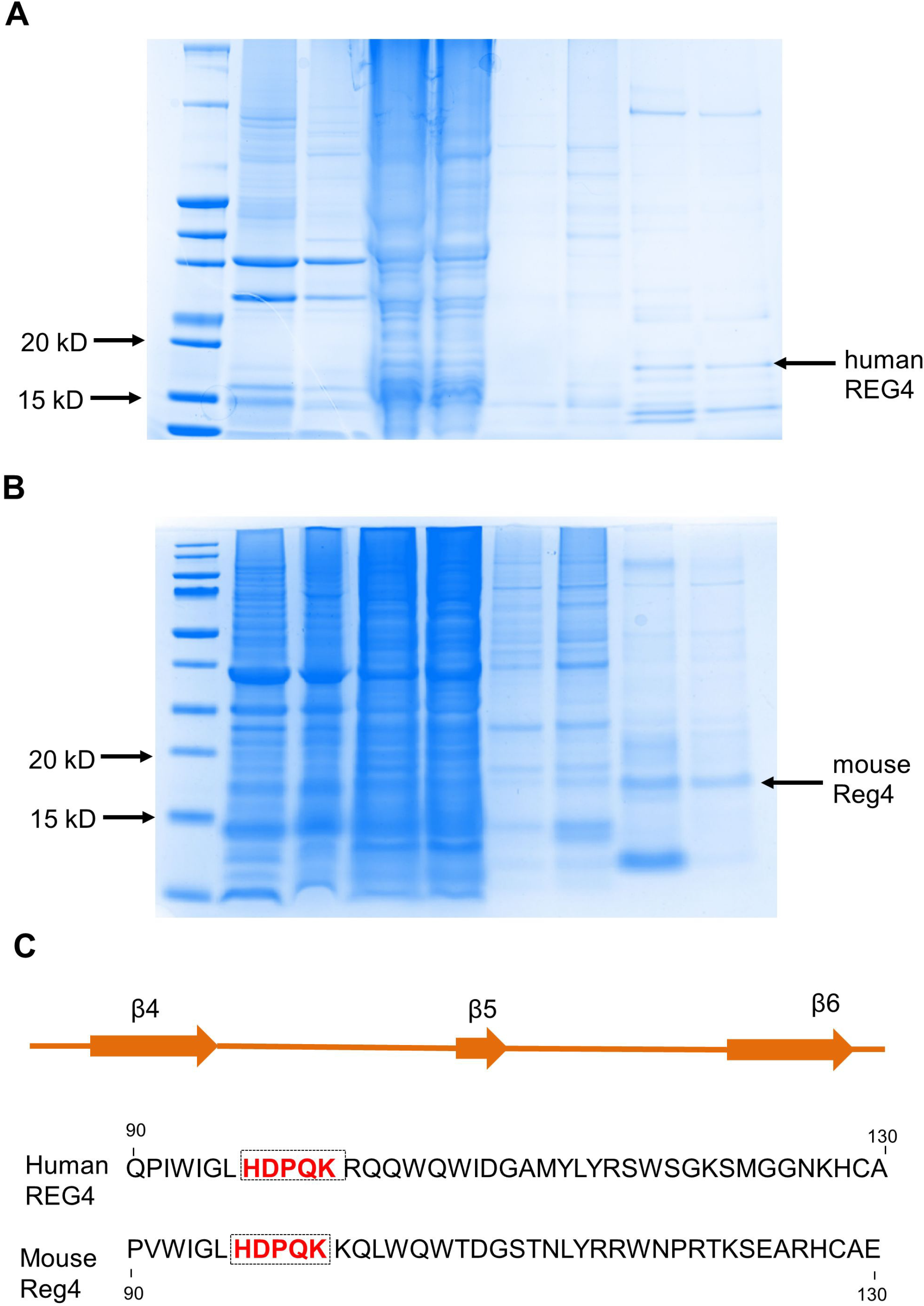
Purified of Recombinant human REG4 and mouse Reg4. **(A)** Coomassie Brilliant Blue staining of human REG4. Black arrows highlight human Reg4. **(B)** Coomassie Brilliant Blue staining of mouse Reg4. Black arrows highlight mouse Reg4. **(C)** Comparing the motif of human REG4 and mouse Reg4. The shared motif “HDPQK” are highlighted in red frames.

**Figure 3.**
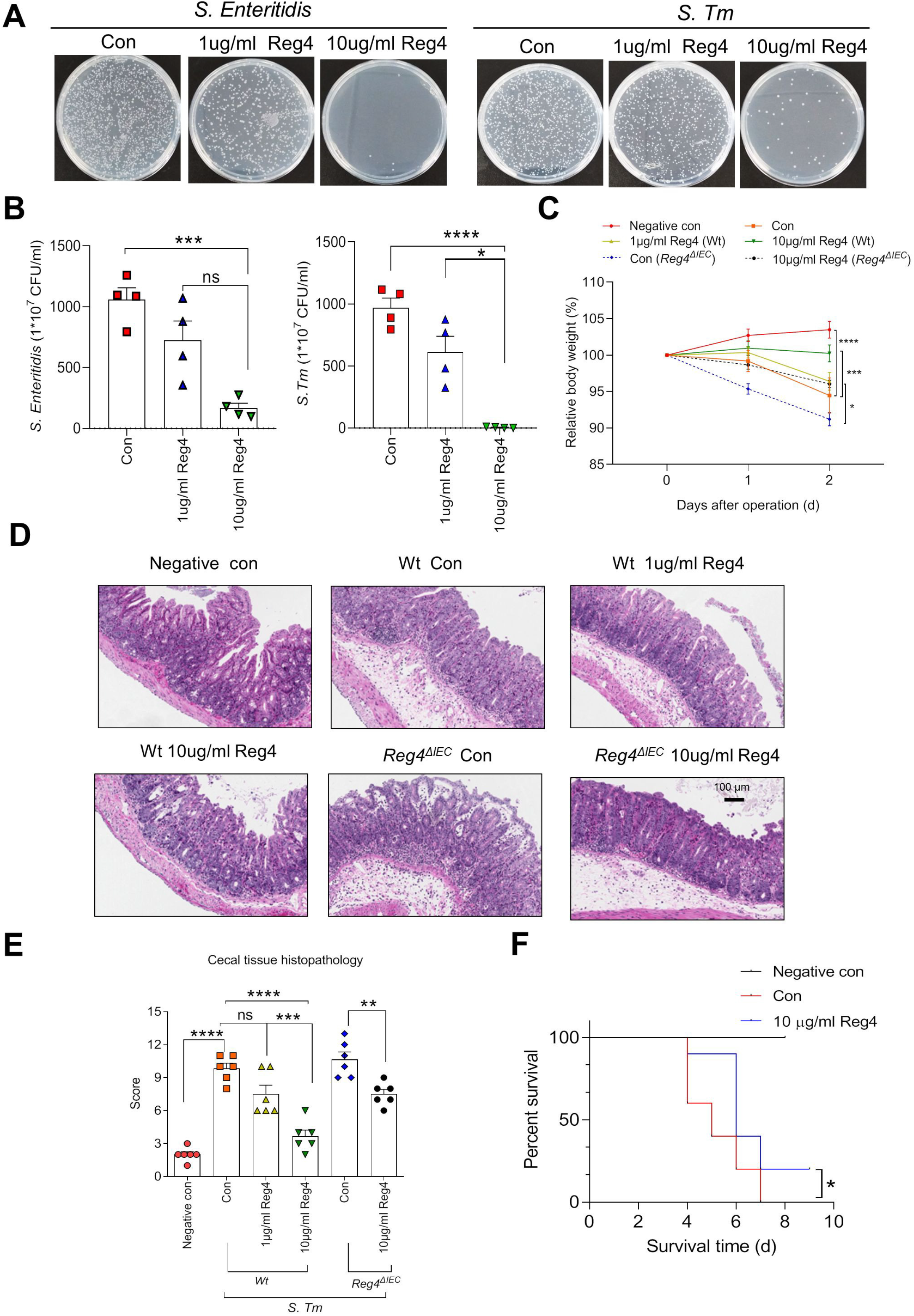
Reg4 has bactericidal activity against *Salmonella* Typhimurium. **(A and B)** The bactericidal activity of mouse Reg4 against *S. Tm* and human REG4 against *S. Enteritidis.* Data are mean ± SEM; unpaired t test was used for statistical analysis. **(C)** Percentage of weight change of mice (n = 10 in wild-type mice, n = 9 in *Reg4 ^ΔIEC^* mice). Data are mean ± SEM; one-way ANOVA was used for statistical analysis. **(D and E)** Histology of mice was assessed by using cecal tissue section, and histological score was plotted. Data are mean ± SEM; one-way ANOVA was used for statistical analysis

**Figure 4.**
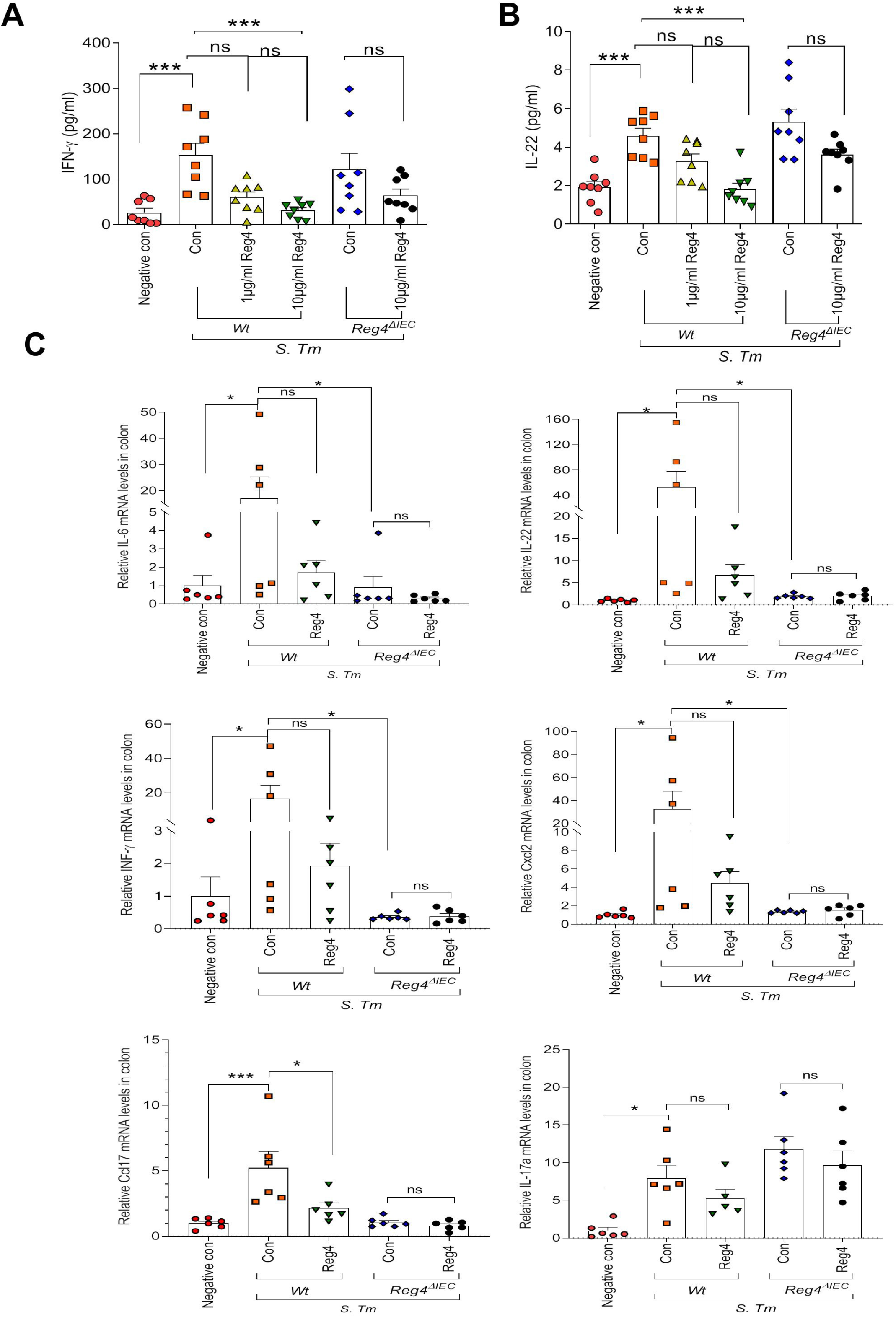
Reg4 ameliorate intestinal inflammation. **(A)** ELISA analysis of serum IL-22 of mice (n = 8 per group). Data are mean ± SEM; one-way ANOVA was used for statistical analysis. **(B)** ELISA analysis of serum INF-γ of mice (n = 8 per group). Data are mean ± SEM; one-way ANOVA was used for statistical analysis. **(C)** Quantitative reverse transcription (RT)–PCR of the expression of proinflammatory cytokine. Values were normalized to Actin expression. Data are mean ± SEM; one-way ANOVA was used for statistical analysis; ns, not significant (*p* ≥ 0.05); * *p* < 0.05; ** *p* < 0.01, *** *p* < 0.005; **** *p* < 0.001.

A previous study has shown that Reg3γ promoted the physical separation of host and microbiota and limited the bacterial colonization of the mucosal surface in the small intestine(13). We predicted that Reg4 may also limit the colonization of *S. Tm*. To investigate the role of Reg4 in intestinal colonization, the numbers of viable Salmonella cells were determined in fresh fecal and cecum samples of infected mice. As expected, the intestinal *S. Tm* colonization was decreased in Reg4-treated mice compared to control mice (Fig. 5A-B). Consistently, IHC demonstrated less *S. Tm* colonized in the cecum in the Reg4-treated mice (Fig. 5C). To further investigate the effects of Reg4 on *S. Tm* translocation, the numbers of CFU in extraintestinal organs were determined. Significantly decreased recovery of viable *S. Tm* from the liver, spleen, and mesenteric lymph node (MLN) of Reg4-treated mice compared to that in control mice (Fig. 6A-C). Besides, the *Reg4^ΔIEC^* mice had more intestinal colonization and severer translocation of S. Tm (Fig. 5A-B and Fig.6A-C). Reg4 protein decreased the intestinal colonization and extraintestinal translocation in *Reg4^ΔIEC^* mice (Fig. 5A-C and Fig.6A-C). Based on these results, it appears that Reg4 exerts beneficial effects on intestinal inflammation during acute Salmonella colitis that is dependent on its antagonism of *S. Tm* colonization.

**Figure 5.**
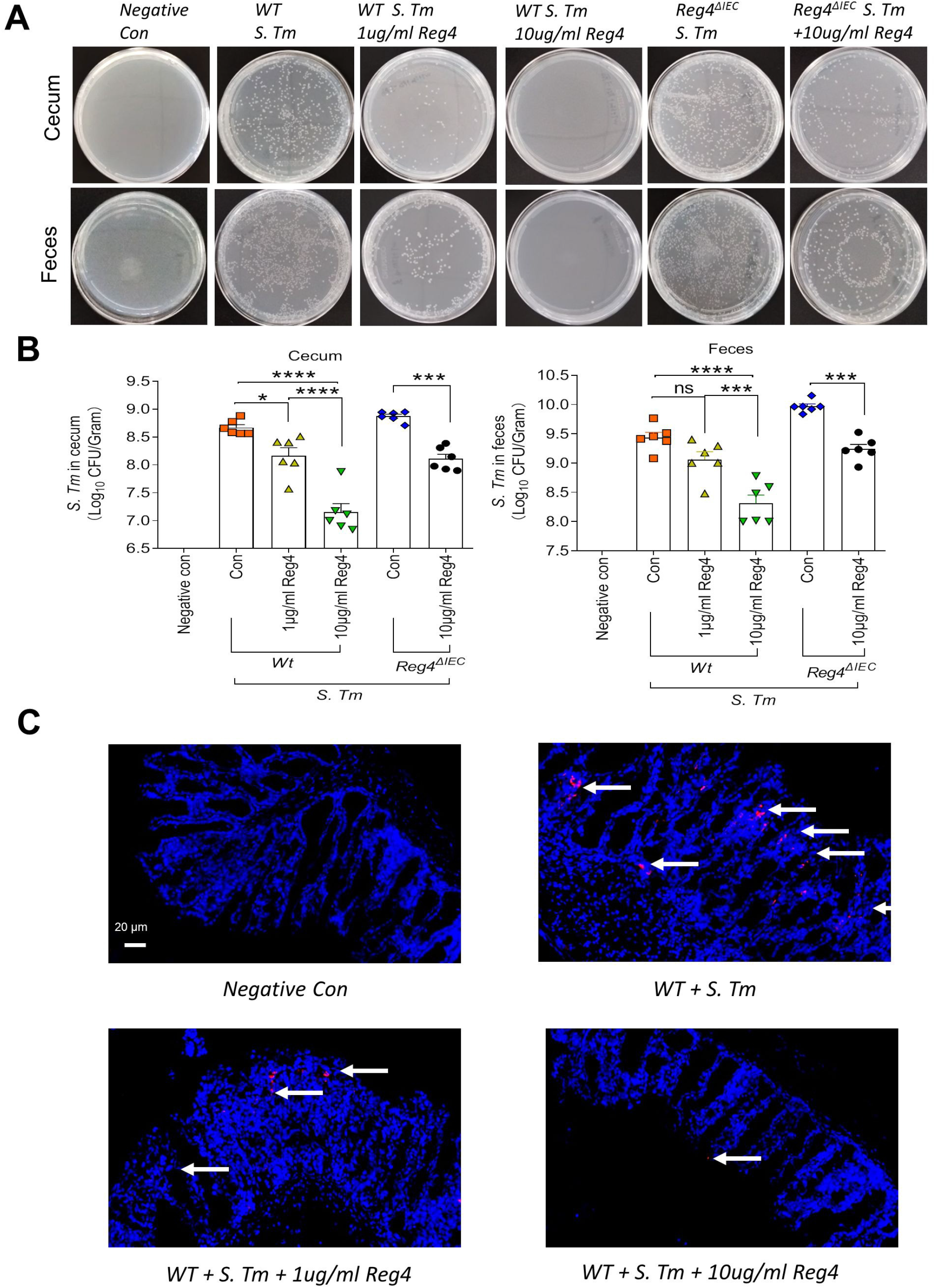
Reg4 decrease the location of *Salmonella* Typhimurium. **(A and B)** *S. Tm* location detected by CFU per gram of tissue on LB plates from cecal tissue and feces (n = 6 per group). Data are mean ± SEM; one-way ANOVA was used for statistical analysis; ns, not significant (p ≥ 0.05); * p < 0.05; ** p < 0.01, *** p < 0.005; **** p < 0.001. **(C)** Visualization of *Salmonella* (red) in mouse cecum detected by IF. White arrows highlight Salmonella.

**Figure 6.**
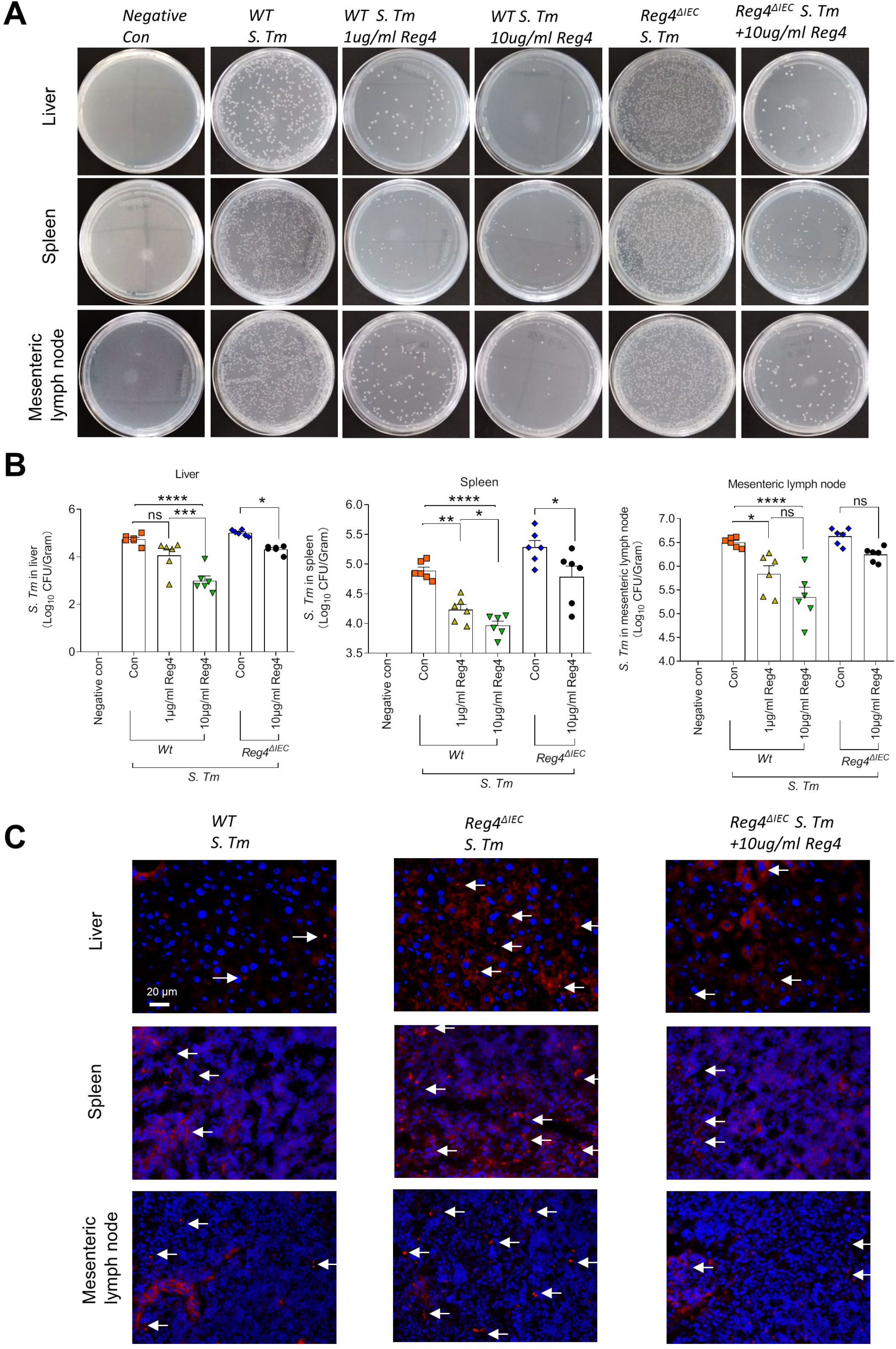
Reg4 decrease the translocation of Salmonella Typhimurium. **(A and B)** *S. Tm* dissemination detected by CFU per gram of tissue on LB plates from liver, spleen and MLN (n = 6 per group). Data are mean ± SEM; one-way ANOVA was used for statistical analysis; ns, not significant (*p* ≥ 0.05); * *p* < 0.05; ** *p* < 0.01, *** *p* < 0.005; **** *p* < 0.001. **(C)** Visualization of *Salmonella* (red) in mouse liver, spleen, and MLN detected by IF. White arrows highlight *Salmonella*.

### The antibacterial effect of Reg4 depend on its “HDPQK” motif

The previous study demonstrated that human REG4 protein contained two functional modules that can support calcium-independent sugar-binding(14). The site I consist of Arg46, Tyr86, Gln87, Arg88, Asn135, Asn136, and Asn137. The site II is composed of His97, Asp98, Gln100, Lys101, Arg102, Gln103, and His128. The site II was shared in human REG4 and mouse Reg4 by comparing their sequences (Fig. 2F). It contains a conserved “HDPQK” motif formed by His, Asp, Pro, Gln, and Lys, which exhibit large chemical shift changes in the mannan titration experiment(14). Based on the fact that both Reg4 and REG4 killed the bacterial *in vitro*, we hypothesized that the antibacterial effect of Reg4 and human REG4 may depend on the “HDPQK” motif. To test it, we first purified the recombinant mutant mouse Reg4 and human REG4 by deleted the “HDPQK” motif. Both mutant mouse Reg4 and human REG4 exhibited no bactericidal activity in *in vitro* growth assay (Fig.7A and 7B). Comparing to those of untreated *S. Tm*-infected mice, weight loss and histological score improved in the wild-type Reg4-treated mice, but not in mutant Reg4-treated mice (Fig.7D and 7E). Moreover, the mutant Reg4 cannot limit the intestinal colonization of *S. Tm* and extraintestinal translocation (Fig.7F and 7G). Taken together, the antibacterial effect of Reg4 and human REG4 mainly depends on its conserved “HDPQK” motif.

**Figure 7.**
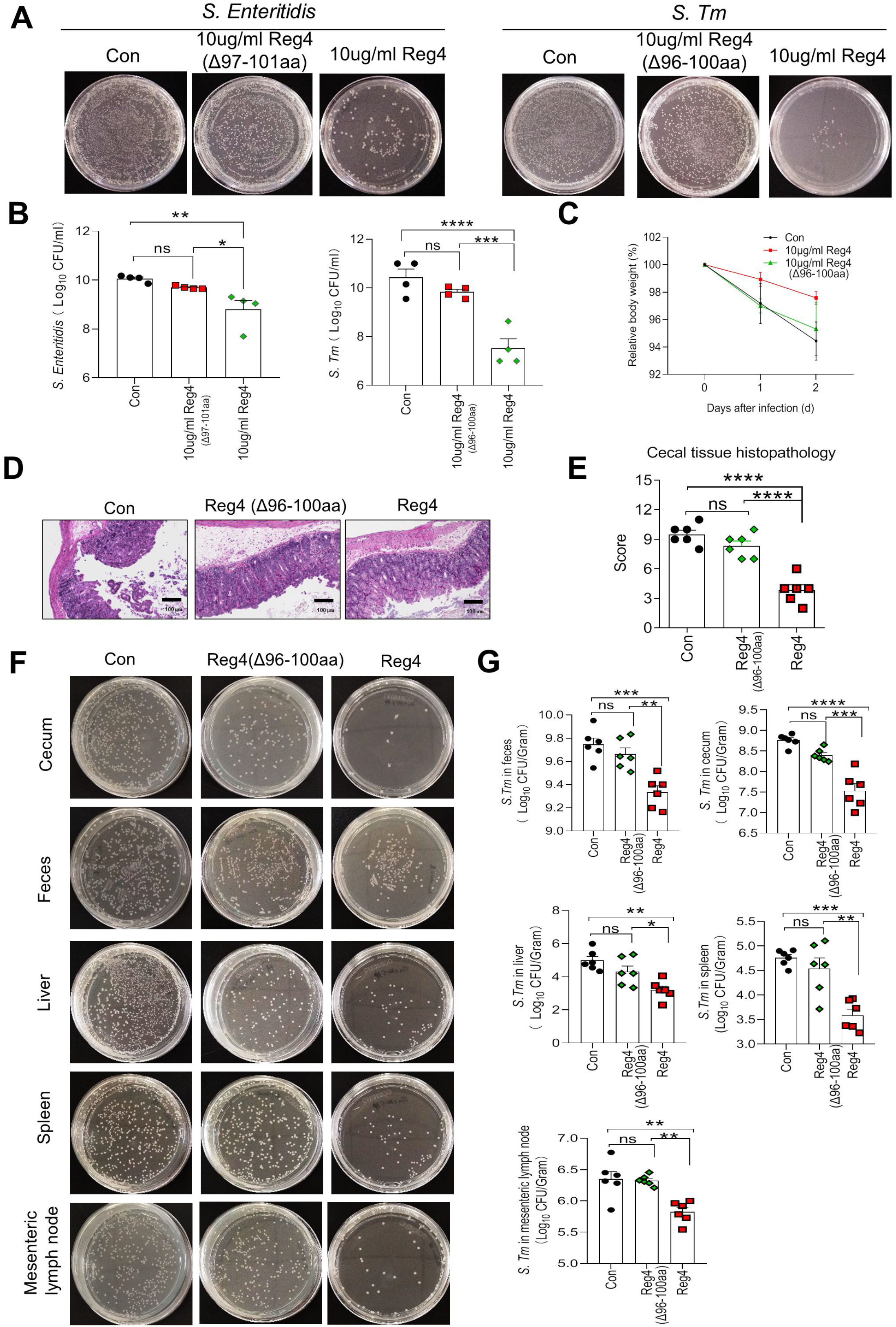
The mutant Reg4 loss bactericidal activity. **(A and B)** The bactericidal activity of mutant Reg4 against *S. Tm* and human REG4 against *S. Enteritidis* (n = 8 per group). Data are mean ± SEM; unpaired t test was used for statistical analysis. **(C)** Percentage of weight change of mice (n = 10 per group) treated with Reg4 or mutant Reg4. Data are mean ± SEM; one-way ANOVA was used for statistical analysis. **(D and E)** Histology of mice (n = 6 per group) treated with mutant Reg4 was assessed by using cecal tissue section, and (E) histological score was plotted. Data are mean ± SEM; one-way ANOVA was used for statistical analysis. **(F and G)** Location and translocation of *S. Tm* in mice (n = 6 per group) treated with mutant Reg4. Data are mean ± SEM; one-way ANOVA was used for statistical analysis; ns, not significant (*p* ≥ 0.05); * *p* < 0.05; ** *p* < 0.01, *** *p* < 0.005; **** *p* < 0.001.

### Reg4 promotes the segregation of *S. Tm* and epithelial cells

We next addressed the mechanisms by which Reg4 inhibits bacterial invasion of the epithelial layer. Firstly, we tested if REG4 decrease the efficient adhesion and invasion of *S. Enteritidis* on epithelial cells *in vitro*. Caco2 cells were incubated with *S. Enteritidis* with or without REG4 treatment, and then *S. Enteritidis* that attached to Caco2 cells were analyzed. The adhesion and invasion of *S. Enteritidis* were markedly diminished in the presence of REG4 (Fig.8A and 8B). We investigated whether Reg4 was able to inhibit bacterial motility. *S. Tm* was co-cultured with Reg4, then added to the center of the semisolid agar, and incubated for 4 h. In the presence of mouse Reg4 protein, the motility of *S. Tm* was reduced (Fig.8C). Similarly, the migratory activity of *S. Enteritidis* was inhibited in the presence of human REG4 (Fig.8D).

**Figure 8.**
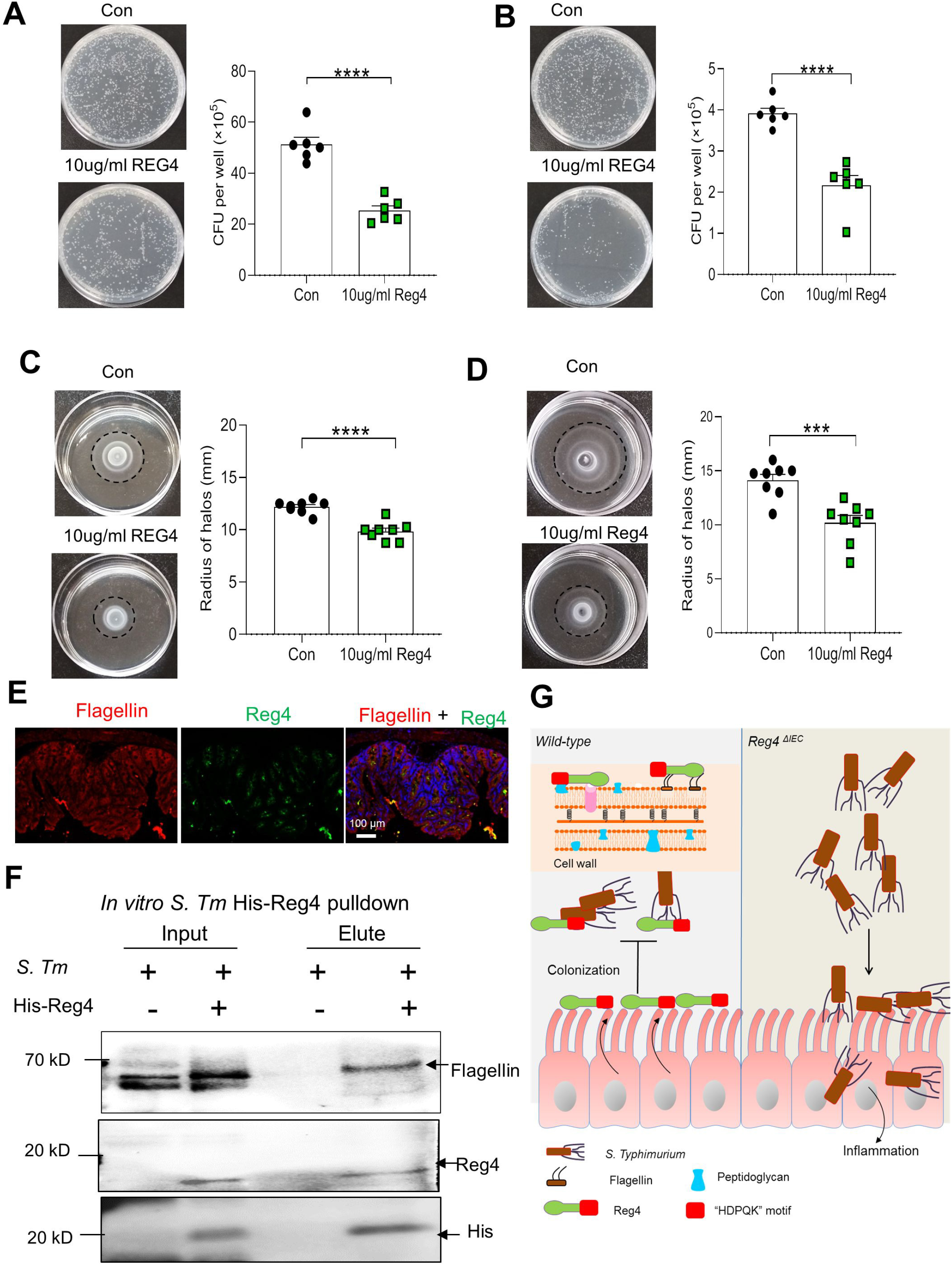
Reg4 reduce motility, adhesion and invasion of *Salmonella* Typhimurium. **(A)** The mean CFU of *S. Enteritidis* adhering to Caco2 cells. Data are mean ± SEM; unpaired t test was used for statistical analysis. **(B)** The mean CFU of *S. Enteritidis* invading to Caco2 cells. Data are mean ± SEM; unpaired t test was used for statistical analysis. **(C)** Motility of *S. Enteritidis* in semisolid agar with or without human REG4. Representative photos and the radii of motility halos at 4 h. Data are mean ± SEM; unpaired t test was used for statistical analysis. **(D)** Motility of *S. Tm* in semisolid agar with or without Reg4. Representative photos and the radii of motility halos at 4 h. Data are mean ± SEM; unpaired t test was used for statistical analysis; ns, not significant (*p* ≥ 0.05); * *p* < 0.05; ** *p* < 0.01, *** *p* < 0.005; **** *p* < 0.001. **(E)** Immunostaining with anti-Reg4 antibody (green), anti-Flagellin (red) and DAPI (blue) of Carnoy’s fixed cecal tissue sections. Scale bars, 100 μ m. **(F)** Reg4/flagellin complexes were formed by incubating the lysate of *S. Tm* with His-tagged Reg4. Binding was assessed by Ni-NTA column pull-down followed by western blotting with anti-Flagllin, anti-Reg4, and anti-His antibodies. **(G)** Graphical Abstract illustrating the role of Reg4 in defensing the intestine against *Salmonella* infection.

Previous studies suggested that flagellin plays a central role in the infection processes(15). We hypothesized that Reg4 binds specifically to bacterial flagellin. Immunofluorescence (IF) assay showed flagellin co-localized with Reg4 in the cecum. To examine whether Reg4 interacts with the flagellin of *S. Tm*, we generated the His-tagged Reg4. Using the lysates of *S. Tm* containing flagellin, we performed pull-downs with recombinant His-tagged Reg4. His-tagged Reg4 was captured by using Ni-NTA column and binding of flagellin was assessed by immunoblotting. Interestingly, the Reg4 was capable of interacting with flagellin (Fig. 8F). To sum up, these findings indicated that Reg4 binds to flagellin, reducing their motility, and thereby inhibits bacterial adhesion and invasion of the epithelial cells.

## Discussion

*Salmonella* Typhimurium is one of the most common causes of food-borne illness and is a major cause of diarrheal diseases, respectively(2, 16). Over the years, the antimicrobial resistance of *Salmonella* Typhimurium is increasing, imposing a serious threat to global health. Therefore, there is an urgent need for alternative strategies that can both kill pathogens and resolve harmful inflammation. As a part of intestinal immunity, AMPs are a promising approach, which are produced to inhibit or directly kill microorganisms and thus maintain gut homeostasis(17, 18). Here, we demonstrate that Reg4 can act as a novel AMP to enhance host defense against *Salmonella* infection and attenuate *Salmonella* Typhimurium-induced intestinal inflammation. Additionally, Reg4 can reduce the location of *Salmonella* Typhimurium in the intestine and translocation in extraintestinal organs. This study reveals that the Reg4 works through a conserved “HDPQK” motif in the C-type lectin-like domain. Reg4 binds the flagella of *Salmonella* Typhimurium and reduces their motility, adhesion, and invasion.

*Salmonella* Typhimurium infection induces a host inflammatory immune response(16). Inflammatory responses regulating proinflammatory cytokine production and gene expression in the intestine, ultimately result in intestinal inflammation(16). Consistent with these previous reports, we found that infection of *S. Tm* induced intestinal inflammation. The *S. Tm* infected mice suffered from more weight loss and lower survival rate. They had more severe intestinal inflammation manifested as higher histological scores of the cecum, and more proinflammatory cytokine production and expression. Many published reports have shown that AMPs play an important role in the defense against pathogenic microorganisms and the regulation of the innate and adaptive immune pathways(18–20). Moreover, AMPs represent a powerful tool against drug-resistant pathogens because they are evolutionarily conserved, with a limited propensity for resistance(17).

Reg family proteins may act as AMPs because they have a C-type lectin-like domain, which recognizes their bacterial targets by binding peptidoglycan carbohydrate and then limit direct contact between bacteria and the intestinal epithelium(21). Previous studies have elucidated the bactericidal activity of REG3G and REG3B(13, 21, 22). REG3G has a bactericidal effect on bacteria Gram-positive bacteria and promotes the spatial segregation of intraluminal bacteria and the intestinal epithelium(13, 21, 23). REG3B has bactericidal activity against Gram-negative bacteria and defenses mice against intestinal infection and location of *Salmonella* Typhimurium(24, 25). However, to date, the bactericidal activity of Reg4, another important member of the Reg family, is little understood. Here, the comprehensive microarray and analysis of mRNA expression profiling in pig ileum tissues and analysis of Reg4 mRNA expression in mouse colon all revealed that Reg4 was increasingly expressed during Salmonella infection. In addition, higher expression of Reg4 is associated with less Salmonella fecal shedding. The lower shedding of *S. Tm* means less inflammation(26). These revealed that Reg4 may be involved in the process of intestinal inflammation induced by *Salmonella*. Then we purified the Reg4 proteins and determined that Reg4 can exert strong bactericidal activity against *Salmonella* Typhimurium *in vitro* as well as *in vivo*. *In vivo* experiments, Reg4 supplements suppressed the growth of *S. Tm* and *S. Enteritidis*. Consistent with these findings, the therapeutic administration of Reg4 for the *S. Tm*-infected mice can attenuate body weight loss and improve survival rate. In addition, Reg4 can ameliorate intestinal inflammation by decreasing both production and gene expression of proinflammatory cytokines. It shows that Reg4 can regulate the excessive production of proinflammatory cytokines that occurs in response to pathogen induction. The study also found that Reg4 can decrease the location of *S. Tm* in the intestine and reduce the number of bacteria that translocate to extraintestinal tissues, such as the liver, spleen, and MLN. The bactericidal activity of Reg4 is also supported by the findings from *Reg4*-deficiency mice. *Reg4* deficiency induced more severe intestinal inflammation, the greater number of *S. Tm* located in the intestine and translocated to extraintestinal tissues. However, administration of Reg4 can reverse these negative impacts.

As depicted above, the C-type lectin-like domain plays an important role in the bactericidal activity of Reg family protein(21–25). The previous study demonstrated that human REG4 protein contained two functional modules that can support calcium-independent sugar-binding(14). Interestingly, by comparing the modules of human REG4 and mouse Reg4 protein, we found the conserved “HDPQK” motif formed by His, Asp, Pro, Gln and Lys, which exhibit large binding ability with mannan(14). Based on the fact that both Reg4 and REG4 killed the bacteria *in vitro*, we hypothesized that the antibacterial effect of Reg4 and human REG4 may depend on the “HDPQK” motif. As expected, the bactericidal activity of Reg4 was decreased when the “HDPQK” motif was deleted. Moreover, mutant Reg4 cannot ameliorate the intestinal inflammation of *S. Tm* infected mice and limit the intestinal colonization and extraintestinal translocation of *S. Tm*. These findings suggest that the antimicrobial effect of Reg4 and human REG4 mainly depends on its conserved “HDPQK” motif.

We next explored the proposed mechanisms by which Reg4 inhibits bacterial invasion of the host. Previous studies demonstrated that Flagella-dependent motility is crucial for Salmonella pathogenesis by enabling directed movement towards host epithelial cells(15, 27). Flagella also play a critical role in other infection processes, including colonization, adhesion, invasion, biofilm formation, and immune system modulation(15, 28, 29). In this study, we found that REG4 can inhibit the adhesion and invasion of *S. Enteritidis* on Caco2 cells. In addition, the migratory activity of *S. Tm* and *S. Enteritidis* in the semisolid agar was also inhibited by Reg4. We thus speculated that Reg4 may bind with flagella and thus exert the bactericidal effect. Given that the main component of flagella is flagellin, we pulled down the lysates of *S. Tm* with recombinant His-tagged Reg4, and detected the presence of flagellin. Indeed, Reg4 was capable of binding with flagellin. Consistent with these findings, immunofluorescence analysis also revealed the co-location of Reg4 and flagellin. These findings indicate that Reg4 binds to flagella, thereby inhibits bacterial motility and decreasing the adhesion and invasion of *Salmonella* Typhimurium in the intestinal lumen.

Together, Reg4 exhibits strong bactericidal activity that protected the mice against Salmonella Typhimurium-induced inflammation, intestinal location, and extra-intestinal translocation. Reg4 may recognize *Salmonella* Typhimurium through the conserved “HDPQK” motif, and then binds with the flagella of *Salmonella* Typhimurium, thereby reduce their motility, adhesion, and invasion to epithelial cells. As many enteric pathogenic bacteria have flagella, Reg4 may be a promising therapeutic drug for treating enteric pathogen-induced intestinal inflammation diseases.

## Materials and Methods

### Bacterial strains and growth

All bacterial strains used in the study are listed in Key Resource Table. Salmonella Typhimurium SL-1344 (*S. Tm*, kanamycin-resistant) and Salmonella enterica subsp. enterica serovar Enteritidis (*S. Enteritidis*) were routinely cultured overnight in Lysogeny broth (LB) (10 g/l tryptone, 5 g/l yeast extract, and 5 g/l NaCl) at 37 °C with shaking at 200 rpm or on LB agar plates. Kanamycin (50 μg/ml) was added when required for strain selection. The number of colony-forming units (CFU) per ml was counted. Strains were stored at −80 °C in glycerol.

### Housing and husbandry of experimental animals

Wild type C57BL/6 mice were purchased from Shanghai Jihui Laboratory Animal Care Co., Ltd (Shanghai, China). The *Reg4 ^flox/flox^* (*Reg4^fl/fl^*) mice and Villin-cre+ mice were bred. They were crossed to generated *Reg4* conditional knockout mice (*Reg4^ΔIEC^*). The genotype of the *Reg4*^fl/fl^, Villin-cre+, and *Reg4^ΔIEC^* loci was determined by PCR assays. All mice used for experiments were age and sex-matched. They were kept under specified pathogen-free conditions in the Department of Model Animal Research, Xinhua Hospital Affiliated to Shanghai Jiao Tong University School of Medicine. All animal experiments in this study were approved by the Animal Research Committee of Xinhua Hospital Affiliated to Shanghai Jiao Tong University School of Medicine.

### Infection of mice with cultured bacteria

Infection experiments were performed as described previously(30). Seven-week-old wild type C57BL/6 mice and *Reg4^ΔIEC^* mice were allocated to control and experimental groups randomly. Mice were pretreated with 20 mg of kanamycin 24 h before infection. Then, they were infected by oral gavage with bacteria (1×10^8^ CFU) containing sterile PBS or varying concentrations of Reg4. Mice were kept in individual cages to avoid transmission between mice. Animals were sacrificed on the 2nd day after infection by cervical dislocation. For chronic infections, mice were infected by gavage with *S. Tm*. The experimental mice were injected with Reg4 at 10 μg/ml and the control mice were injected with sterile PBS daily post-infection. Changes in weight and survival were closely monitored by daily observation.

### Colonization of mice with cultured bacteria

For analysis of the bacterial burden, fresh fecal pellets, cecum, the right lobe of the liver, the mesenteric lymph nodes, and the spleens were isolated and suspended in 1ml sterile PBS using a Tissue grinding device, serially diluted, and plated on LB plates containing the appropriate antibiotics for CFU determination. CFU counts were performed by two researchers in a blinded manner. Bacterial burden was measured as follows: tissues were harvested and homogenized at 1 mg per ml equivalent in LB, serially diluted 1:10 in LB, and 100 μl was plated on LB agar plates. For *S. Tm* CFU determination, LB agar plates were supplemented with kanamycin. After 16 h, the number of CFU was counted.

### Production of recombinant reg4 protein

The recombinant protein was generated as previously described(31, 32). Briefly, the coding sequence of DNA, including mouse Reg4 (NM_026328-6His), mutant mouse Reg4 (NM_026328(del96-100aa)-6His), human REG4 (NM_001159352-6His), and mutant human REG4 (NM_001159352(del97-101aa)-6His), was cloned into the pET-28 Expression vector (Shanghai Genechem Co., Ltd) with an N-terminal 6-His tag and transformed into BL21(DE3) competent cells. Expression of the recombinant protein was induced by the addition of 1 mM isopropyl-β-thiogalactopyranoside (IPTG) once the optical density at 600 nm (OD_600_) reached 0.6. The induced culture was further incubated at 37 °C and the cells were harvested by centrifugation. Harvested cells were resuspended in lysis buffer and disrupted with lysozyme. The soluble fraction was clarified by centrifugation and the supernatant was run through a Ni-NTA column. Then the column was washed and eluted to get the eluted protein. The eluted protein was buffered exchanged into endotoxin-free phosphate buffered saline (PBS) using a sterile PD MidiTrap G-25 column (GE Healthcare). Finally, the protein was concentrated to 1 mg/ml using a vivaspin 20 (GE Healthcare) and store at −80 °C. The quality of the purified protein was analyzed by SDS-PAGE and analyzed by Coomassie Brilliant Blue staining.

### Microarray data

The GEO database (https://www.ncbi.nlm.nih.gov/geo/) stores original submitter-supplied records as well as curated datasets. The gene expression dataset GSE27000 and GSE23063 was obtained from the GEO database. This dataset had been generated using the platform GPL3533 [Porcine] Affymetrix Porcine Genome Array.

### *In vitro* growth assays

*S. Tm* and *S. Enteritidis* (2×10^5^ CFU/ml) were incubated in LB medium (2 ml) with the indicated concentrations of the recombinant Reg4 protein for 24 h at 37 °C. The bacterial cultures were applied to LB agar plates and incubated at 37 °C for 16 h, and the number of CFU per ml was calculated.

### Histopathology

Cecal tissue were fixed in 4% formaldehyde and stained with hematoxylin and eosin. The pathology of cecal samples was scored blindly by two independent pathologists according to the established scoring scheme(30). Each section was evaluated for the presence of submucosal edema, polymorphonuclear granulocytes infiltrate, surface erosions, and decrease of goblet cells. These scores were added together to generate the overall pathological score for each sample. It ranges from 0 to 13 and covers varying degree of inflammation as follow: 0 - 2 = normal, 3 - 4 = mild inflammation, 5 - 8 = moderate inflammation, 9 - 13 = severe inflammation.

### Immunohistochemistry

Immunohistochemistry (IHC) was performed to assess the distribution of *S. Tm* and Reg4 in organs. Paraffin-embedded sections were dewaxed, hydrated, and blocked with 1% bovine serum albumin (BSA) in PBS. It was stained with the primary antibody of Reg4 antibody or Anti-Flagellin antibody in an optimal concentration (Reg4 antibody, dilution 1:100; Anti-Flagellin antibody, dilution 1:200). After washed with PBS, the slides were incubated with secondary antibodies conjugated to fluorescein isothiocyanate. The nuclei were counterstained with 4′, 6-diamidino-2-phenylindole (DAPI). The positive staining signals were detected with fluorescence microscopy.

### Enzyme-linked immunosorbent assay

The concentrations of cytokines, including IL-22 (Cat # YCJL29108) and IFN-γ (Cat # D721025-0096), were measured in the serum by using enzyme-linked immunosorbent assay (ELISA) kits according to the manufacturer’s instructions.

### Gene expression by Quantitative RT-PCR

Colon tissue was homogenized and total RNA was isolated with the RNeasy Mini kit following the protocol of the manufacture (Qiagen). The level of the genes was detected using the High-Capacity cDNA Reverse Transcription kit and the SYBR-Green Universal Master Mix kit. The list of the real-time PCR primers is provided in the Table S1.

### Pull-down assay of Reg4 binding to flagella

The assay of Reg4 biding to flagella was performed according to the previously described protocol(33). Bacteria were grown overnight at 37 °C. Then, the bacteria were suspended in lysis buffer and destroyed with lysozyme. The supernatant was collected by centrifugation, and incubated with the Reg4 protein at 4 °C overnight to form the Reg4 / flagellin complexes. Binding was assessed by Ni-NTA-agarose pull-down followed by western blotting with anti-Flagellin, REG4 antibody, and Penta·His Antibody.

### Motility assay of bacteria in semisolid agar

Bacteria were cultured in LB medium at 37 °C until OD_600_ reached 0.6. Then, the bacteria were mixed with Reg4 (10 μg/ml) or PBS. The mixture was applied on the semisolid LB agar (0.3%) plates. Motility was assessed by measuring the radius of circles formed by bacterial migration at 4 h after the bacterial inoculation.

### Bacterial adhesion and invasion assay on Caco-2 cells

The Caco-2 cells were obtained from the Chinese Academy of Sciences (Shanghai, China) and cultured in Dulbecco’s modified Eagle’s medium (DMEM; Cat # 11965-092; Gibco, Grand Island, NY) supplemented with 20% heat-inactivated fetal bovine serum (FBS; Cat # 26140079; Gibco) in a humidified atmosphere with 5% CO_2_ at 37 °C. The bacterial adhesion and invasion assay with modified were performed as described previously(27, 33). Briefly, Caco-2 cells (2.5 × 10^5^ cells per ml) were seeded in 24-well plates and incubated with 5% CO_2_ at 37 °C. *S. Enteritidis* was grown to mid-log phase, diluted in sterile PBS, and incubated with or without Reg4 protein at 10 μg/ml. Then, Caco-2 cells were infected with *S. Enteritidis* for 2 h in the humidified atmosphere with 5% CO_2_ at 37 °C. The contact of the bacteria with the Caco-2 cells was forced by centrifugation at 500 × g for 5 min. Afterward, the Caco-2 cells were washed with 1 × PBS extensively to remove unbound bacteria and lysed using 1% Triton X-100. The lysate was serially diluted and plated on LB agar plates to calculate the bacterial CFU per ml. For the invasion assay, the Caco-2 cells were incubated with 100 ug/ml gentamycin for 2 h to kill the external bacteria. Afterward, the Caco-2 cells were washed and lysed, and bacterial CFU per ml was determined as described above.

### Quantification and statistical analysis

Experimental results are presented as means ± standard error of mean and analyzed by GraphPad Prism 8 Software (GraphPad, San Diego, CA). the difference between the control and experimental groups was evaluated using a two-tailed unpaired Student’s t-test, one-way ANOVA, and two-way ANOVA, appropriately. Kaplan-Meier survival curves were performed. A *P*-value < 0.05 was regarded as statistical significance.

## Acknowledgements

We appreciate the support by Jing Zhu, Yaying You and Hui Cai in the experimental operation.

## Data Availability

Raw data were generated at Xinhua hospital. Derived data supporting the findings of this study are available from the corresponding author Yongtao Xiao on request.

## References

1. Fabrega A, Vila J. 2013. Salmonella enterica serovar Typhimurium skills to succeed in the host: virulence and regulation. Clin Microbiol Rev 26:308–41.

2. Crump JA, Sjolund-Karlsson M, Gordon MA, Parry CM. 2015. Epidemiology, Clinical Presentation, Laboratory Diagnosis, Antimicrobial Resistance, and Antimicrobial Management of Invasive Salmonella Infections. Clin Microbiol Rev 28:901–37.

3. Michael GB, Schwarz S. 2016. Antimicrobial resistance in zoonotic nontyphoidal Salmonella: an alarming trend? Clin Microbiol Infect 22:968–974.

4. Ouellette AJ. 2010. Paneth cells and innate mucosal immunity. Curr Opin Gastroenterol 26:547–53.

5. Chen Z, Downing S, Tzanakakis ES. 2019. Four Decades After the Discovery of Regenerating Islet-Derived (Reg) Proteins: Current Understanding and Challenges. Front Cell Dev Biol 7:235.

6. Hartupee JC, Zhang H, Bonaldo MF, Soares MB, Dieckgraefe BK. 2001. Isolation and characterization of a cDNA encoding a novel member of the human regenerating protein family: Reg IV. Biochim Biophys Acta 1518:287–93.

7. Grun D, Lyubimova A, Kester L, Wiebrands K, Basak O, Sasaki N, Clevers H, van Oudenaarden A. 2015. Single-cell messenger RNA sequencing reveals rare intestinal cell types. Nature 525:251–5.

8. van Beelen Granlund A, Ostvik AE, Brenna O, Torp SH, Gustafsson BI, Sandvik AK. 2013. REG gene expression in inflamed and healthy colon mucosa explored by in situ hybridisation. Cell Tissue Res 352:639–46.

9. Oue N, Mitani Y, Aung PP, Sakakura C, Takeshima Y, Kaneko M, Noguchi T, Nakayama H, Yasui W. 2005. Expression and localization of Reg IV in human neoplastic and non-neoplastic tissues: Reg IV expression is associated with intestinal and neuroendocrine differentiation in gastric adenocarcinoma. J Pathol 207:185–98.

10. Nanakin A, Fukui H, Fujii S, Sekikawa A, Kanda N, Hisatsune H, Seno H, Konda Y, Fujimori T, Chiba T. 2007. Expression of the REG IV gene in ulcerative colitis. Lab Invest 87:304–14.

11. Tsuchida C, Sakuramoto-Tsuchida S, Taked M, Itaya-Hironaka A, Yamauchi A, Misu M, Shobatake R, Uchiyama T, Makino M, Pujol-Autonell I, Vives-Pi M, Ohbayashi C, Takasawa S. 2017. Expression of REG family genes in human inflammatory bowel diseases and its regulation. Biochem Biophys Rep 12:198–205.

12. Xiao Y, Lu Y, Wang Y, Yan W, Cai W. 2019. Deficiency in intestinal epithelial Reg4 ameliorates intestinal inflammation and alters the colonic bacterial composition. Mucosal Immunol 12:919–929.

13. Vaishnava S, Yamamoto M, Severson KM, Ruhn KA, Yu X, Koren O, Ley R, Wakeland EK, Hooper LV. 2011. The antibacterial lectin RegIIIgamma promotes the spatial segregation of microbiota and host in the intestine. Science 334:255–8.

14. Ho MR, Lou YC, Wei SY, Luo SC, Lin WC, Lyu PC, Chen C. 2010. Human RegIV protein adopts a typical C-type lectin fold but binds mannan with two calcium-independent sites. J Mol Biol 402:682–95.

15. Chaban B, Hughes HV, Beeby M. 2015. The flagellum in bacterial pathogens: For motility and a whole lot more. Semin Cell Dev Biol 46:91–103.

16. Gart EV, Suchodolski JS, Welsh TH, Jr., Alaniz RC, Randel RD, Lawhon SD. 2016. Salmonella Typhimurium and Multidirectional Communication in the Gut. Front Microbiol 7:1827.

17. Browne K, Chakraborty S, Chen R, Willcox MD, Black DS, Walsh WR, Kumar N. 2020. A New Era of Antibiotics: The Clinical Potential of Antimicrobial Peptides. Int J Mol Sci 21.

18. Mookherjee N, Anderson MA, Haagsman HP, Davidson DJ. 2020. Antimicrobial host defence peptides: functions and clinical potential. Nat Rev Drug Discov 19:311–332.

19. Zhang LJ, Gallo RL. 2016. Antimicrobial peptides. Curr Biol 26:R14–9.

20. Diamond G, Beckloff N, Weinberg A, Kisich KO. 2009. The roles of antimicrobial peptides in innate host defense. Curr Pharm Des 15:2377–92.

21. Mukherjee S, Zheng H, Derebe MG, Callenberg KM, Partch CL, Rollins D, Propheter DC, Rizo J, Grabe M, Jiang QX, Hooper LV. 2014. Antibacterial membrane attack by a pore-forming intestinal C-type lectin. Nature 505:103–7.

22. Wang L, Fouts DE, Starkel P, Hartmann P, Chen P, Llorente C, DePew J, Moncera K, Ho SB, Brenner DA, Hooper LV, Schnabl B. 2016. Intestinal REG3 Lectins Protect against Alcoholic Steatohepatitis by Reducing Mucosa-Associated Microbiota and Preventing Bacterial Translocation. Cell Host Microbe 19:227–39.

23. Cash HL, Whitham CV, Behrendt CL, Hooper LV. 2006. Symbiotic bacteria direct expression of an intestinal bactericidal lectin. Science 313:1126–30.

24. van Ampting MT, Loonen LM, Schonewille AJ, Konings I, Vink C, Iovanna J, Chamaillard M, Dekker J, van der Meer R, Wells JM, Bovee-Oudenhoven IM. 2012. Intestinally secreted C-type lectin Reg3b attenuates salmonellosis but not listeriosis in mice. Infect Immun 80:1115–20.

25. Miki T, Holst O, Hardt WD. 2012. The bactericidal activity of the C-type lectin RegIIIbeta against Gram-negative bacteria involves binding to lipid A. J Biol Chem 287:34844–55.

26. Knetter SM, Bearson SM, Huang TH, Kurkiewicz D, Schroyen M, Nettleton D, Berman D, Cohen V, Lunney JK, Ramer-Tait AE, Wannemuehler MJ, Tuggle CK. 2015. Salmonella enterica serovar Typhimurium-infected pigs with different shedding levels exhibit distinct clinical, peripheral cytokine and transcriptomic immune response phenotypes. Innate Immun 21:227–41.

27. Horstmann JA, Lunelli M, Cazzola H, Heidemann J, Kuhne C, Steffen P, Szefs S, Rossi C, Lokareddy RK, Wang C, Lemaire L, Hughes KT, Uetrecht C, Schluter H, Grassl GA, Stradal TEB, Rossez Y, Kolbe M, Erhardt M. 2020. Methylation of Salmonella Typhimurium flagella promotes bacterial adhesion and host cell invasion. Nat Commun 11:2013.

28. Rossez Y, Wolfson EB, Holmes A, Gally DL, Holden NJ. 2015. Bacterial flagella: twist and stick, or dodge across the kingdoms. PLoS Pathog 11:e1004483.

29. Duan Q, Zhou M, Zhu L, Zhu G. 2013. Flagella and bacterial pathogenicity. J Basic Microbiol 53:1–8.

30. Barthel M, Hapfelmeier S, Quintanilla-Martinez L, Kremer M, Rohde M, Hogardt M, Pfeffer K, Russmann H, Hardt WD. 2003. Pretreatment of mice with streptomycin provides a Salmonella enterica serovar Typhimurium colitis model that allows analysis of both pathogen and host. Infect Immun 71:2839–58.

31. Jarret A, Jackson R, Duizer C, Healy ME, Zhao J, Rone JM, Bielecki P, Sefik E, Roulis M, Rice T, Sivanathan KN, Zhou T, Solis AG, Honcharova-Biletska H, Velez K, Hartner S, Low JS, Qu R, de Zoete MR, Palm NW, Ring AM, Weber A, Moor AE, Kluger Y, Nowarski R, Flavell RA. 2020. Enteric Nervous System-Derived IL-18 Orchestrates Mucosal Barrier Immunity. Cell 180:50–63 e12.

32. Ho MR, Lou YC, Lin WC, Lyu PC, Huang WN, Chen C. 2006. Human pancreatitis-associated protein forms fibrillar aggregates with a native-like conformation. J Biol Chem 281:33566–76.

33. Okumura R, Kurakawa T, Nakano T, Kayama H, Kinoshita M, Motooka D, Gotoh K, Kimura T, Kamiyama N, Kusu T, Ueda Y, Wu H, Iijima H, Barman S, Osawa H, Matsuno H, Nishimura J, Ohba Y, Nakamura S, Iida T, Yamamoto M, Umemoto E, Sano K, Takeda K. 2016. Lypd8 promotes the segregation of flagellated microbiota and colonic epithelia. Nature 532:117–21.

